# Effects of Drought and Salinity, *etc.* on Root Pressure of *Dracaena sanderiana*

**DOI:** 10.1101/2024.05.14.594061

**Authors:** Weizhi Zhang, Jinfeng Song, Ying Sang, Min Zhang, Jiaqi Zheng, Yuyang Liu

## Abstract

**Aim:** Root pressure plays a crucial role in maintaining tissue hydration and ensuring the functionality of the plant’s hydraulic system while facilitating cellular and tissue growth. The study focuses on investigating the impact of drought, salinity, *etc.* on root pressure of *Dracaena sanderiana*.

**Methods:** Pressure sensors (PX26-100GV) were used to measure the diurnal variations in root pressure of soil-cultured and hydroponic *D. sanderiana* with heights ranging from 60 to 80 cm. The study also examined various influencing factors such as drought stress, salinity stress, temperatures, mineral concentration, change in leaves and roots that affect root pressure. This approach provides valuable insights into the role of root pressure in mediating water upward movement in xylem.

**Essential findings:** The research showed: (1) Both soil-cultured and hydroponic *D. sanderiana* exhibited positive root pressure throughout the day, demonstrating apparent diurnal pattern of high pressures at the day and low pressures at night. The maximum root pressure measured in the experiments was 96 kPa; (2) For soil-cultured *D. sanderiana*, root pressure decreased under drought and salinization (NaCl), while the diurnal rhythm remained unchanged; (3) For hydroponic *D. sanderiana*, under simulated drought (PEG 6000) and salt stress (NaCl), the rate of decrease in root pressure accelerated with increasing concentrations of PEG 6000 and NaCl, displaying diurnal rhythm at low concentrations but losing this pattern at high concentrations; (4) Under room temperature, root pressure of hydroponic *D. sanderiana* decreased with lower water temperature and increased with higher temperatures, but did not increase at 40℃; low concentrations of nitrate (KNO_3_) generally increased root pressure, however, root pressure decreased when concentration reached 4 g·L^-1^, and the diurnal rhythm remained unaffected; (5) For hydroponic *D. sanderiana*, the presence of leaves on the stem below the root pressure measurement point resulted in significantly lower root pressure compared to plants without leaves on the stem, while still maintaining a regular diurnal rhythm. Removal of all root hairs and the entire root system led to a rapid decline in root pressure to negative values, followed by a slow recovery to around 0 kPa. In conclusion, relative to its height, the root pressure of *D. sanderiana* is sufficient to transport water required for transpiration to the top of the plant, effectively countering the impact and function of transpirational pull. Moreover, root pressure shows a stable diurnal rhythm of higher values during the day and lower at night. Environmental factors such as drought, salinity stress, temperatures, mineral concentration, and changes in leaves and roots significantly affect root pressure.

## Introduction

Root pressure, a hydrostatic force generated in plant roots, first identified by Hales in 1727, has regained attention as a critical physiological process (Knipfer *et al*., 2015). Root pressure can help explain phenomena such as nocturnal sap flow in tall trees, embolism repair, budburst before spring leafing, flowering, and recovery in fire-affected standing trees. Root pressure is evident in maize (*Zea mays L.*) and grapevines (*Vitis vinifera L.*) (Yang *et al*., 2012; Gleason *et al*., 2017) and essential for embolism repair during seasonal shifts in certain trees and lianas (Hölttä *et al*., 2018). Additionally, root pressure plays a crucial role in re-establishing xylem hydration following mild drought stress, representing a common hydrological adaptation in bamboo and certain herbaceous plants. (Andersen & Brodbeck, 1989; Clearwater *et al*., 2007; Cobb *et al*., 2007; Yin *et al*., 2018). Further investigation into positive xylem pressure, including its generation sites within the roots and stems, is warranted across a broader range of plant species (Schenk *et al*., 2021).

The origin of root pressure remains a contentious subject despite the various proposed theories (Fan *et al*., 1994). Currently, the most accepted explanations are the osmotic theory, metabolic theory, and the hypothesis of water’s upward co-transport (Zhai *et al*., 2011; Zhang *et al*., 2022). The osmotic theory posits roots as osmotic pumps, during the night, when transpiration is minimal or absent, living cells in the conductive tissue transport mineral ions and soluble organics into the lumens., thereby altering the luminal water potential and inducing a continuous water influx. Concurrently, soil water, driven by the potential gradient, permeates the xylem via root hairs, cortex, and endodermis, creating a positive hydrostatic force (Dong, 2003; Singh, 2016). Zholkevich and colleagues proposed the metabolic theory, which posits an osmotic gradient between the internal water flow in roots, external soil solutions, and the exudates released by root systems. (Zholkevich, 1981<u>_EN-</u> <u>REF_1</u>). Root pressure generated in the xylem lumens due to solute osmotic potential is associated with the difference in water potential across the endodermis and the physiological metabolism of the root system. The adenosine triphosphate (ATP) necessary for this active water transport is provided by plant respiratory metabolism, underscoring the critical role of respiration in root water absorption processes (Wilson & Kramer, 1949). The hypothesis of upward co-transport of water, introduced by Wegner (2013) based on the co-transport model (Zeuthen, 2010), integrates osmotic and metabolic theories. It suggests that aquaporins (AQPs) facilitate water movement within thin-walled cells in the xylem, actively driving upward water transport in the presence of free energy gradients of ions and soluble sugars. This hypothesis emphasizes the consideration of water and solute demand of plant roots when elucidating the mechanism of root pressure generation (Wegner, 2014; Singh, 2016; Zhang *et al*., 2022).

Methods for measuring root pressure include the U-tube manometer. (Grossenbacher, 1938), external pressure application (White, 1938), bubble pressure gauges (Fisher, *et al*., 1997), and pressure sensor techniques (Wang *et al*., 2015;Miller, 1985; Cao *et al*. 2012). The use of a pressure sensor (pressure transducer) is widespread due to its ability to rapidly equilibrate and continuously monitor root pressure in real-time (Henzler *et al*., 1999; McClure & Franklintong, 2010). Variations in root pressure among different species are well-documented. White (1938) observed that root-to-xylem water secretion persisted unabated even when outer pressure exceeded 0.6 MPa in hydroponic five-year-old tomato plants (*Solanum lycopersicum*). Grossenbacher (1938) connected U-tube mercury manometer to main stem of hydroponic plants for 35 days, noting root pressures often exceeding the 0.15 MPa measurement range of the instrument (Grossenbacher, 1938). In corn, root pressures of 0.42 MPa was recorded using pressure sensors, sometimes reaching up to 0.6 MPa (Miller, 1985). Cao *et al*. (2012) utilized pressure sensors to measure root pressure in bamboo during rainy season, discovering a positive correlation between root pressure and bamboo height, with a maximum root pressure of 0.2 MPa observed in 20 m tall bamboo. Woody lianas also exhibit root pressure, ranging from 2–138 kPa, and maintain positive pressure throughout the day(Wang, 2015). Distinct diurnal patterns are observed in plant bleeding and root pressure, as evidenced in six-year-old walnut trees (*Juglans regia*), where bleeding is higher between 13:00–19:00, decreasing significantly from 19:00 to the following noon in autumn, with less pronounced daily variations in spring (Ding & Xi, 1991). Both soil-cultivated and hydroponic poplar saplings (84K poplar) exhibited a precise diurnal rhythm in detached root systems, with higher root pressure during the day and lower at night (Guo & Wan, 2017). White (1938) further noted that daytime water secretion by the tomato root system elevated the water column above ground level but slowed or halted at night, demonstrating a diurnal cycle even in a small section of the root system (White, 1938; Wang, 2015).

Root pressure is influenced by multiple factors, predominantly relying on the osmotic differential between the endodermis and exodermis. Consequently, any factor impacting transport proteins indirectly affects the presence and magnitude of root pressure (Fan *et al*., 1994; Hachez *et al*., 2006). Water is one of the most critical elements influencing plant root pressure. Barrios-Masias *et al* (2015) found that drought treatment reduces the osmotic potential and hydraulic conductivity of fine roots, leading to earlier root xylem embolization and higher sap osmotic potential, resulting in more significant root pressure upon rehydration(Barrios-Masias *et al*., 2015). Previous studies have indicated an immediate increase in root pressure of *Rhipidocladum racemiflorum* following precipitation (Siligan *et al*., 2017). However, recent research suggests minimal influence of rainfall on root pressure (Wang, 2015). Soil mineral elements also play a significant role in influencing root pressure. Ewers *et al*. (2001) observed that a decrease in soil nitrate concentration led to reduced xylem pressure of walnut, which increased after 4 hours of nitrate concentration enhancement. However, the limiting effect of a certain nitrate concentration level on root pressure remains unknown. NaCl can lower soil water potential, creating osmotic stress and reducing soil water availability, making it difficult for plant roots to absorb water, potentially causing plant water stress or even cellular dehydration (Wang *et al*., 2006; Liu *et al*., 2022; Chen *et al*., 2023). The toxic effects of NaCl might impact root pressure. Temperature is a key limiting factor for tree growth in northern forests (Tryon & Chapin, 1983; Bonan, 1992), affecting the magnitude and diurnal rhythm of root pressure, with maximum root pressure values and root respiration rates decreasing alongside soil temperature (Guo & Wan, 2017). Walnut root pressure is closely related to temperature, exhibiting a simple positive correlation with soil temperature during autumn and spring, a pattern also observed in other tree species exhibiting spring sap flow (Cochard, H. *et al*., 1994; Ewers *et al*., 2001). The presence or absence of leaves also impacts root pressure. In early spring, when air and soil are relatively warm and moist, significant sap flow and root pressure are evident before leaf unfolding (Singh, 2016). After leaf expansion, there is a rapid movement of water within the plant, making it impossible to detect any sap flow or root pressure (Hales, 1727).

*Dracaena sanderiana Sander* (lucky bamboo), an ornamental plant from *Asparagaceae* family, resembles bamboo and can be cultivated in soil and water, thriving well in clear water. Monocot plants like bamboos exhibit notable root pressure, with consistent diurnal cycles of positive pressure at night and negative during the day, representing plants that share a diurnal cycle of hydraulic strategy (Cochard, H *et al*., 1994). Zhang *et al*. 2023 found that hydroponic *D. sanderiana* has positive root pressure all day which is sufficient to support its stature needs, making it an ideal plant for root pressure studies (Zhang, *et al*., 2023). This study measures the root pressure of soil-cultured and hydroponic *D. sanderiana*, examining its diurnal rhythm and the impact of drought, NaCl, temperature, nitrates, leaf presence, and root removal on root pressure. The research aims to provide a theoretical basis for water management in cultivating *D. sanderiana* and offers an ideal plant model for root pressure studies. The response of root pressure to factors in real time and quickly makes it a progressive future to be a method to study plant physiology that plant response to stresses or nutrients and to monitor the plant activity.

## Materials and Methods

### Materials and environment

*Dracaena sanderiana* Sander was cultivated both in soil and water under natural light. Soil-cultured *D. sanderiana* was planted in pots measuring 47 × 24 × 17.5 cm. Peat soil was used with two plants in each pot. Hydroponic *D. sanderiana* was grown in tap water with depth of 10–15 cm. Plants with slender stems (diameter 5.43–7.20 mm) and heights ranging from 60 to 80 cm were selected for the study. Root pressure measurement points were positioned at a height of 27.2–45.2 cm above the ground, with three replicates for each treatment. Experiments were conducted from 2022 to 2023 at west-facing laboratory of the main building of Northeast Forestry University, Harbin, China (45°43′24.6″N, 126°38′21.12″E). During summer, indoor temperatures peaked at 32°C and dropped to a minimum of 22°C; while in winter, the temperature was maintained above 18°C with humidity around 40-60%.

### Root pressure measurement method

*D. sanderiana* was connected with pressure sensor following the procedure of Yang (2012). *D. sanderiana* with thin stem was selected. The upper part of the stem bearing leaves was trimmed and the lower part was connected to pressure sensor (PX26-100GV, OMEGA, USA) using a rubber tube (inter diameter 3 mm) filled with degassed water. Datalogger (CR1000, Campbell Scientific, USA) was connected to pressure sensors to collect data at 15-second intervals and log average values every 15 minutes. Additionally, the datalogger is capable of measuring battery voltage and room temperature. Pressure sensor PX26-100DV, previously used by other researchers, produced consistent values with those obtained using PX26-100GV when its low-pressure end is open to atmosphere. The pressure sensor output is in millivolts (mV) and requires conversions to kilopascals (kPa). PX26-100GV has a range of 0–100 PSI (1 PSI = 6.89 kPa) and, when excited by a 10 V voltage, produces a full-scale output of 100 mV. The corresponding full-scale output voltage (Vmax) for different excitation voltages (Vexc) is calculated as Vmax = Vexc × 100/10, with the maximum pressure being 100 PSI. Therefore, root pressure (kPa) is determined using the formula: 100 PSI/Vmax ×output voltage ×6.89.

### Effects of drought and salinity on soil-cultured D. sanderiana

In drought experiment for soil-cultured *D. sanderiana*, soil was thoroughly watered and then allowed to dry. Root pressure was continuously monitored during this period. In salinization experiment, 200 mmol·L^-1^ NaCl solution was added into pots, saturating the soil thoroughly. Root pressure was recorded before and after NaCl treatment.

### Treatment of drought, salinity, temperature, KNO_3_, leaf presence, and root cutting on hydroponic D. sanderiana

Simulated drought for hydroponically grown *D. sanderiana* was induced by adding polyethylene glycol (PEG) 6000 to solutions with concentrations of 5% (-0.100 MPa), 10% (- 0.200 MPa), and 20% (-0.587 MPa). Salt stress (NaCl) was exposed by immersing plants in NaCl solutions with concentrations of 1, 10, 100, and 200 mmol·L^-1^. For the 200 mmol·L^-1^ NaCl solution, after the root pressure stabilization, the plants were transfered to clear water to assess whether the root pressure could be restored to its pre-stress state. Root pressure was measured before and after the treatments.

For temperature treatment, *D. sanderiana* was transferred from room temperature water to water with temperatures of 0, 15, 35, and 40°C respectively. Subsequently, water temperature gradually returned to room temperature. For KNO_3_ treatment, following a phase of root pressure measurement in clear water, KNO_3_ was added to achieve concentrations of 0.5, 1.0, 2.0, and 4.0 g·L^-1^ in the water. After three days, the plants at 4.0 g·L^-1^ concentration were returned to clear water. Root pressure was continuously monitored before and after the treatments.

Typically, the stem section connected to a pressure sensor does not bear leaves, ensuring a smooth surface free of damage to prevent water loss through leaves, stem breaches, or imperfect connections. For leaf cutting treatment, the pressure sensor was attached to the *D. sanderiana* stem that bears leaves. The leaves were progressively removed, with one-third detached at each day. For root cutting treatment, sterilized scissors were used to sever all fibrous roots or entire root system of *D. sanderiana*. Root pressure was continuously monitored throughout the process.

### Data analysis

Data were organized and processed with Excel 2010. Graphical presentation was conducted using Origin 2018 (OriginLab, Northampton, MA, USA).

## Results

### Effects of drought and NaCl on root pressure of soil-cultured D. sanderiana

The soil-cultured *D. sanderiana* plants exposed to drought displayed distinct diurnal root pressure patterns, ranging from 18 to 70 kPa, with higher pressures during the day and lower pressures at night (Fig. 1). Root pressure remained relatively stable in the initial three days following water cessation. On the second day after water cessation, peak root pressures were observed at 20:45, 18:30 and 18:30, reaching values of 65.23, 64.80 and 50.53 kPa respectively; whereas the lowest values occurred at 10:30, 10:30 and 3:15 with readings of 59.54, 61.60 and 44.76 kPa (Fig. lA). From the fourth day onwards, the root pressure gradually declined for all three plants as drought persisted, resulting in decreased peak values (50.53, 67.23 and 68.23 kPa) and trough values (18.53, 48.52 and 35.05 kPa) on the sixth day, respectively.

**Fig. 1.**
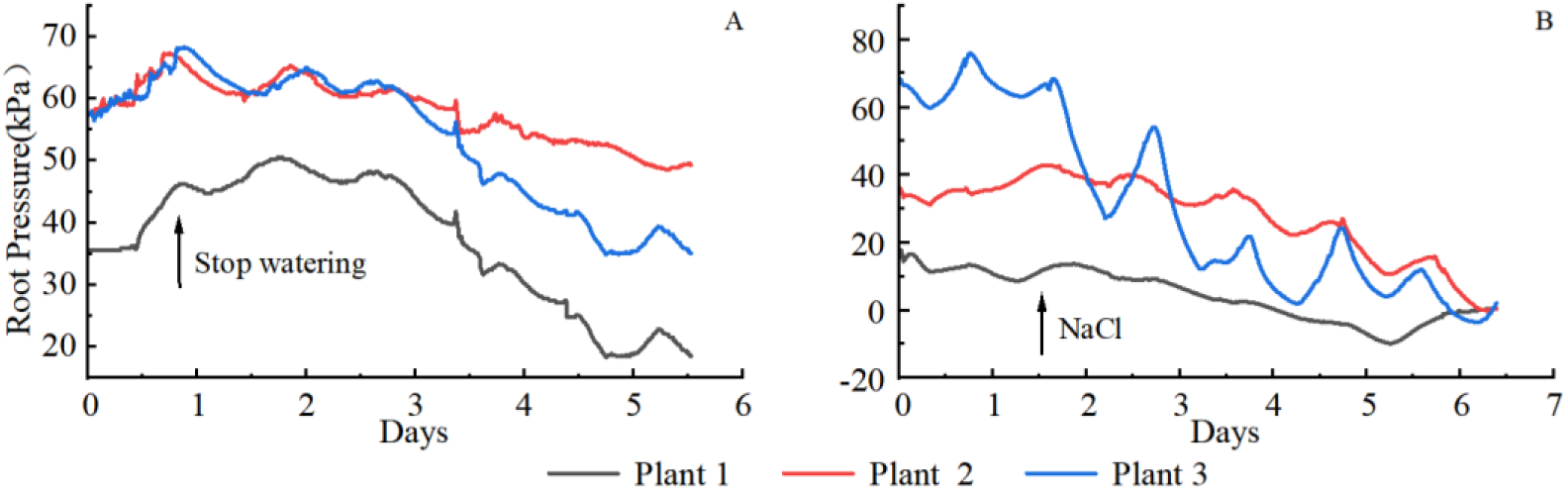
Root pressure of soil-cultured *D. sanderiana* under drought and salt stress.

Under NaCl stress, the three plants exhibited a gradual decrease in root pressure throughout the day while maintaining a distinct diurnal rhythm. Although Plant 1 did not display a precise diurnal rhythm, the rate of decline during the daytime was lower compared to that at night. The peak root pressures decreased from 9.22, 39.83 and 54.05 kPa before treatment to 9.66, 36.50 and 27.11 kPa after six days of treatment (Fig. 1B), indicating a consistent trend of reduction under salt stress while preserving the diurnal rhythm.

### Effects of drought and NaCl on root pressure of hydroponic D. sanderiana

After immersing *D. sanderiana* in PEG-6000 solution, a consistent decline in root pressure was observed across all treatments, with the rate of decline accelerating at higher concentrations. In the 5% PEG-6000 solution, root pressure decreased slowly while maintaining diurnal rhythm variations; after three days, root pressures decreased from pre-treatment values of 67.62, 48.45, 27.20 kPa to -37.43, -0.59, -6.75 kPa respectively and stabilized near these lower values while continuing to exhibit regular diurnal rhythm changes. In the 10% PEG-6000 solution, root pressure dropped from initial values of 27.90, 65.90, 39.47 kPa to 9.24, -11.65, 18.22 kPa after five days and the diurnal rhythm became less pronounced. In the case of the highest concentration tested (20% PEG-6000), root pressure decreased rapidly from initial values of 45.32, 7.17, 13.79 kPa to -42.16, -14.79, -2.76 kPa within seven hours, and then began to rise to stable values around 2.07, -1.02, and 12.16 kPa with minimal fluctuations afterward.

**Fig. 2.**
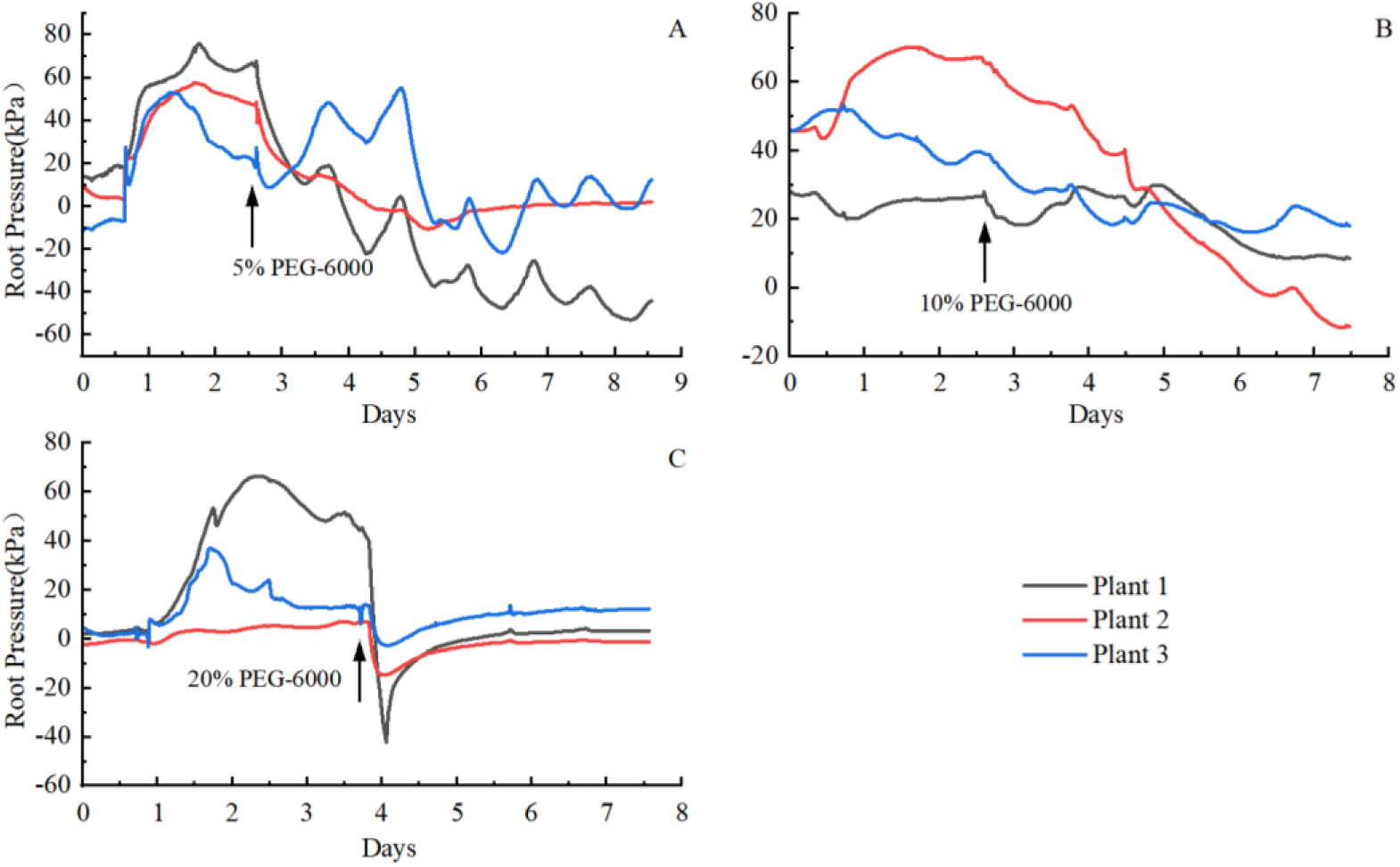
Root pressure of *D. sanderiana* under different concentrations of PEG-6000.

After immersing *D. sanderiana* to NaCl solutions of various concentrations, salt stress consistently resulted in a reduction of root pressure, with higher concentrations leading to more pronounced decreases. Plants in 1 and 10 mmol·L^-1^ solutions exhibited similar patterns, with a rapid increase within a day followed by a gradual decline. In 1 mmol·L^-1^ solution, root pressures surged from 10.43, 51.37, 55.45 kPa to 60.04, 84.03, 79.38 kPa within one day; then slowly decreased to settle around -5.15, 0, and 16.00 kPa after six days while fluctuating near these levels thereafter. In 10 mmol·L^-1^ NaCl solution, root pressures rapidly increased from initial values of 43.16, 28.16, 2.58 kPa to peak values of 60.16, 83.80, 79.21 kPa (Fig. 3A), then gradually declined, eventually stabilizing with minor fluctuations around 2.15, -2.49, 5.69 kPa (Fig. 3B). However, plants in 100 and 200 mmol·L^-1^ NaCl solutions experienced direct decreases in root pressure. In the 100 mmol·L^-1^ solution, Plant 1 dropped from 55.05 kPa to 37.82 kPa within seven hours, then rose and fluctuated around 52.00 kPa for three days before declining. Plants 2 and 3 experienced a rapid decrease from 49.08, 51.83 kPa to -8.89, 7.06 kPa after 20 hours (Fig. 3C).In the 200 mmol·L^-1^ solution, root pressures of all three plants quickly dropped from 78.58, 20.83, 45.98 kPa to 0 kPa within 45 minutes, continuing to drop to their minimum values of - 37.05, -31.62, -47.41 kPa within 115 minutes, then slowly rose back to 0 over 22 hours and remained constant (Fig. 3D). After transferring *D. sanderiana* from 200 mmol·L^-1^ NaCl solutions back to clear water, a slow recovery of root pressures to normal levels was observed for Plants 1 and 3, while no change was observed for Plant 2, indicating the damage inflicted by the 200 mmol·L^-1^ NaCl solution.

**Fig. 3.**
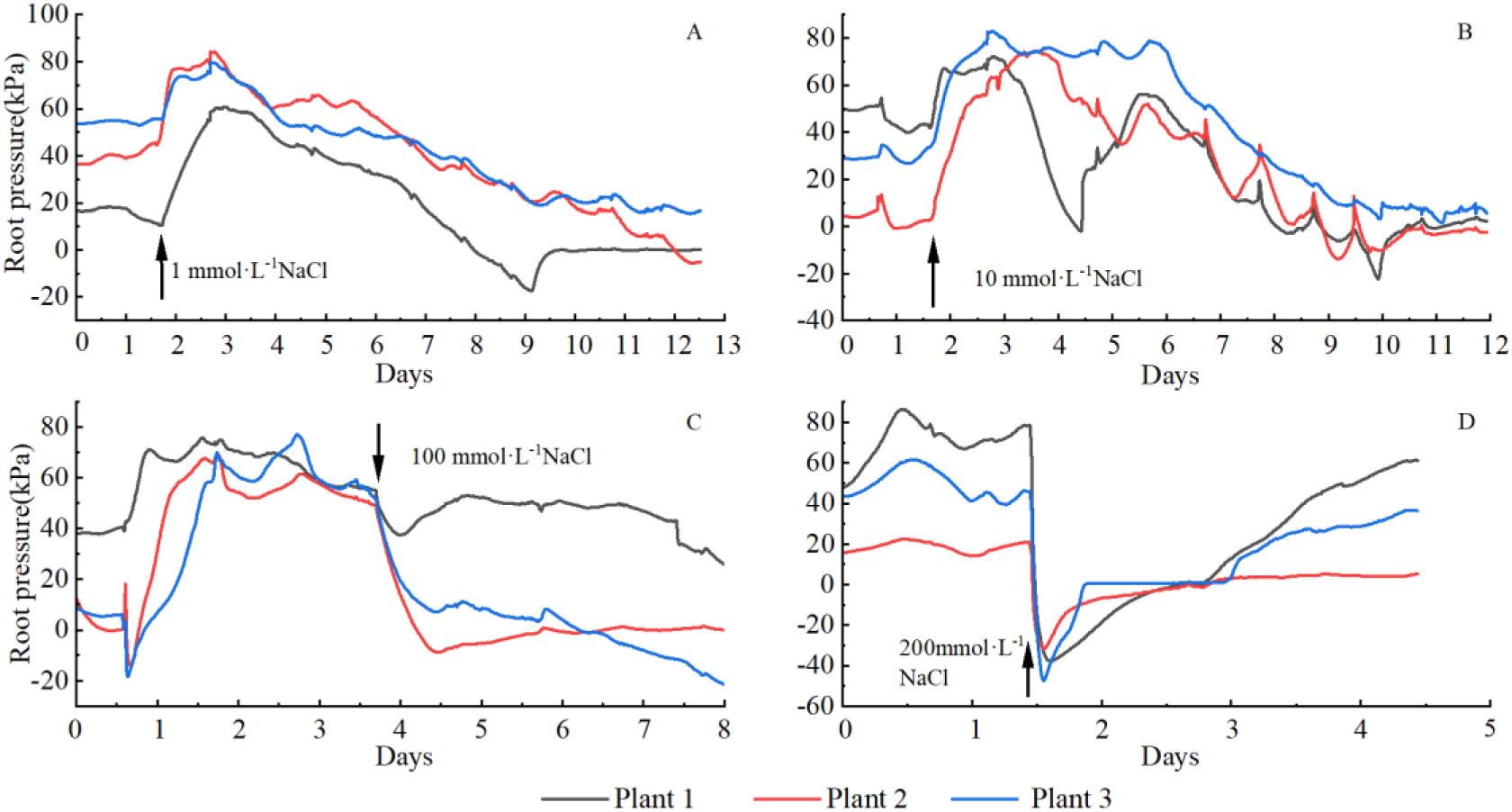
Root pressure of *D. sanderiana* under different concentrations of NaCl.

### Effects of temperature and KNO_3_ on root pressure of hydroponic Dracaena sanderiana

Upon transferring *D. sanderiana* from room-temperature water to 0°C and 15°C water, root pressures initially experienced a rapid linear decline, followed by a slower decrease, eventually rising gradually as the water temperature increased, returning to pre-treatment levels. After immersion in 0°C ice water, root pressures quickly fell, reaching their lowest values after 9.5 hours, decreasing from 37.50, 66.02 and 71.43 kPa to 4.86, 21.09 and 22.77 kPa, respectively. As the water temperature rose again, root pressures also increased to 40.16, 63.68, and 62.52 kPa, showing negligible difference from initial values (Fig. 4A). At a controlled temperature of 15°C, root pressures reached their lowest values after 2 hours, dropping from 50.61, 55.36, 61.00 kPa to 43.91, 46.59, 52.33 kPa, a relatively minor decline. Subsequently, as the water temperature gradually returned to room temperature, root pressures increased to 53.10, 53.10, and 67.36 kPa (Fig. 4B). In contrast, in the case of exposure to 35°C water, root pressure sharply increased in a linear fashion to peak values, increasing from 22.04, 28.69, 31.04 kPa to 53.79, 62.74, 66.71 kPa, then slowly decreased back to 22.16, 21.30, 17.15 kPa as the water temperature gradually returned to room temperature (Fig. 4C). In 40°C water, root pressure quickly dropped, reaching minimum values of 4.63, 4.24, 5.30 kPa after 16 hours (Fig. 4D), subsequently fluctuating diurnally within a range lower than prior levels. Low-temperature treatments resulted in reduced root pressure and altered the diurnal rhythm pattern, while high-temperature treatments altered the diurnal rhythm of root pressure, making the cycle less distinguishable, with the effect and variance increasing at higher temperatures. Overall, the impact of temperature on root pressure is both rapid and immediate.

**Fig. 4.**
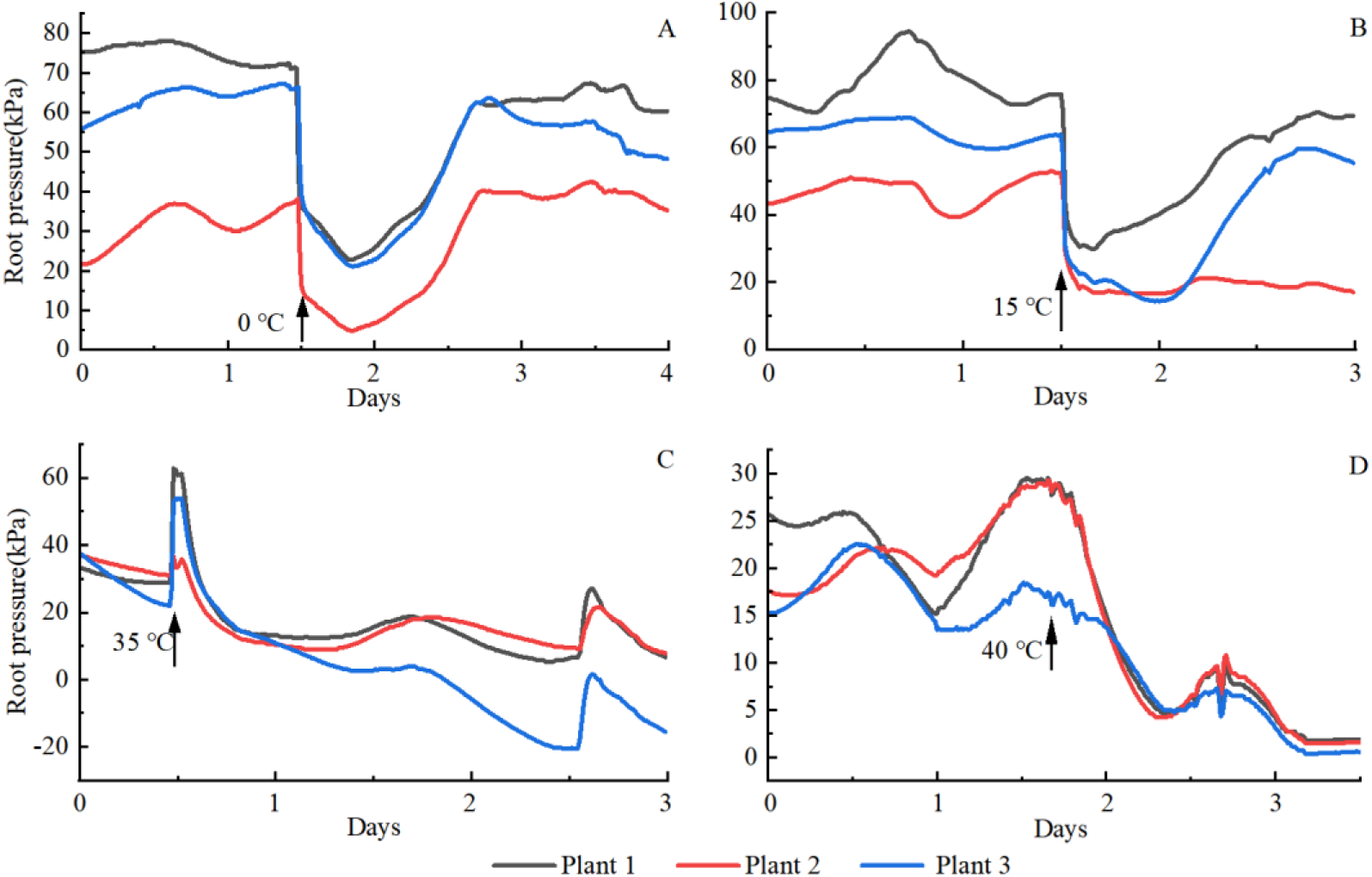
Root pressure of *D. sanderiana* under different temperature.

The impact of varying concentrations of nitrate on root pressure is diverse. In 0.5 g·L^-1^ KNO_3_ solution, root pressure initially decreased rapidly from -0.54, 51.78, 69.06 kPa to -21.73, 21.76, 31.94 kPa, followed by a slow ascent to peak values of 30.01, 46.29, 71.70 kPa (Fig. 5A). In 1.0 g·L^-1^ KNO_3_ solution, root pressure exhibited slight fluctuations before steadily rising to peak values of 33.12, 45.56, 86.01 kPa (Fig. 5B). At a concentration of 2.0 g·L^-1^ KNO_3_, two plants showed minor fluctuations in root pressure, followed by a steady increase, while the peak pressure of the third plant increased significantly, displaying periodic variations (Fig. 5C). In 4.0 g·L^-1^ KNO_3_ solution, root pressure in all three plants rapidly declined; one plant showed a slight increase during the second day, while the other two continued to drop to -19.28, -32.64 kPa. Upon being transferred back to clear water, plant 1 and 2 rose and stabilized within the range of 10-30 kPa. In contrast, plant 3 lost vitality, failing to recover its root pressure (Fig. 5D). This indicates that within a specific range, higher nitrate concentrations lead to more significant increases in root pressure, accompanied by changes in its diurnal rhythm and periodicity. However, once the concentration exceeds a certain threshold, root pressure declines.

**Fig. 5.**
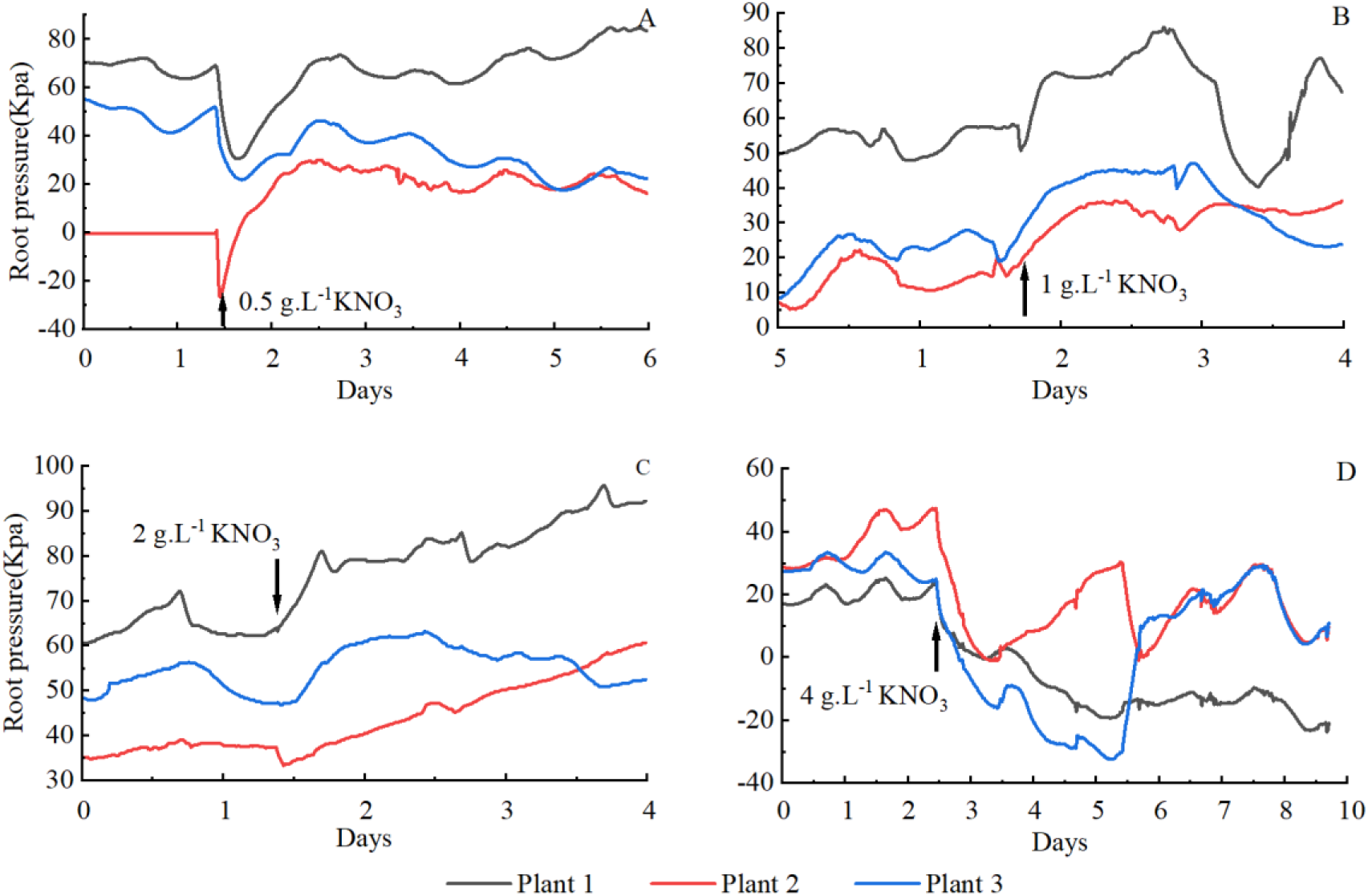
Root pressure of *D. sanderiana* under different concentrations of KNO3.

### Effect of leaves pruning and root cutting on root pressure of hydroponic D. sanderiana

In the presence of leaves on the stem of *D. sanderiana*, root pressure was significantly low, with values 2.37, 3.25, 3.37 kPa, which were only one-tenth to one-twentieth of that in plants without leaves on the stem below the measurement point, also exhibiting apparent diurnal rhythm. On the first day, peak root pressures were recorded at 16:00, with values of 3.97, 3.96 and 3.00 kPa, respectively; while the lowest values appeared at 18:45, 18:30 and 18:15, with corresponding readings of 3.55 kPa, 3.61 kPa and 2.63 kPa. The peaks and troughs occurred successively, with less than a 3-hour difference between them. On the second day, after one third of the leaves were picked off, peak values were observed at 16:15, with values of 3.85, 4.08 and 3.26 kPa, which were a little higher than before. Root pressure increased after leaf removal, though the diurnal rhythm remained unchanged. Removal of leaves resulted in a slight increase in root pressure, with overall higher values post-removal compared to pre-removal, with peak and trough timings similar before and after defoliation.

**Fig. 6.**
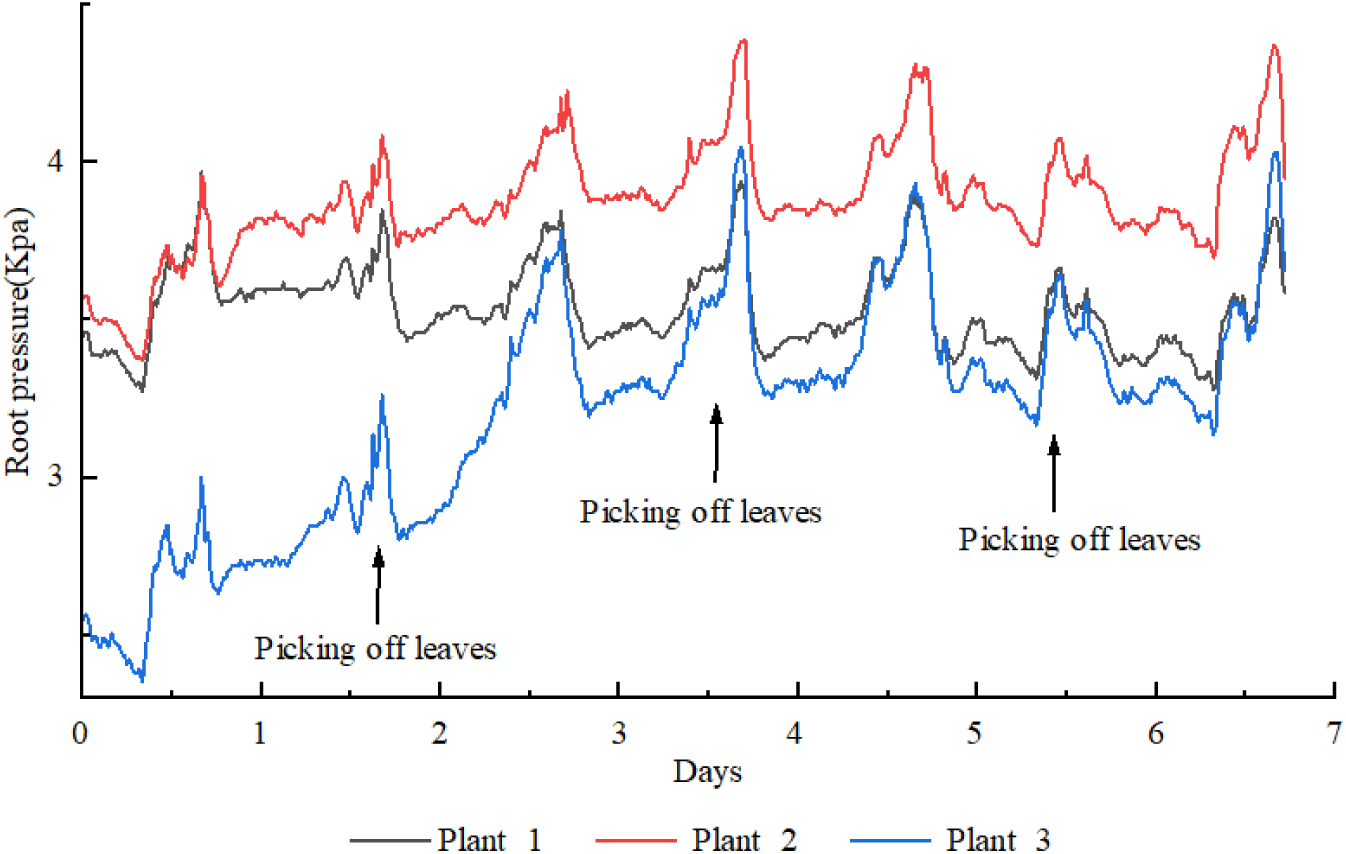
Effect of leaf picking on root pressure of *D. sanderiana*.

Before cutting the fibrous roots, root pressure were relatively high, with peak values of 15.15, 22.52 and 27.17 kPa. After all fibrous roots were severed, the root pressures of Plants 1 and 2 rapidly dropped to 0 kPa within half an hour. Subsequently, plant 1 fluctuated around 0 kPa while Plant 2 continued to decrease further, eventually dropping to -27.22 kPa after 29 hours before slowly recovering back to approximately 0 kPa. In contrast, plant 3 exhibited a slower decline, reaching 0 kPa after 8 hours, followed by a further drop to -12.96 kPa over the next 32 hours before stabilizing with fluctuations around 0 kPa (Fig. 7A). Following the complete removal of roots, root pressure rapidly decreased, taking 9, 12, and 6 hours for the root pressure to fall from 49.36, 38.35, 14.59 kPa to 0 kPa, respectively. The pressure continued to drop to its lowest values before returning to fluctuate around 0 kPa (Fig. 7B). The removal of the root system resulted in the complete loss of positive pressure in *D. sanderiana* along with the cessation of the periodic diurnal rhythm.

**Fig. 7.**
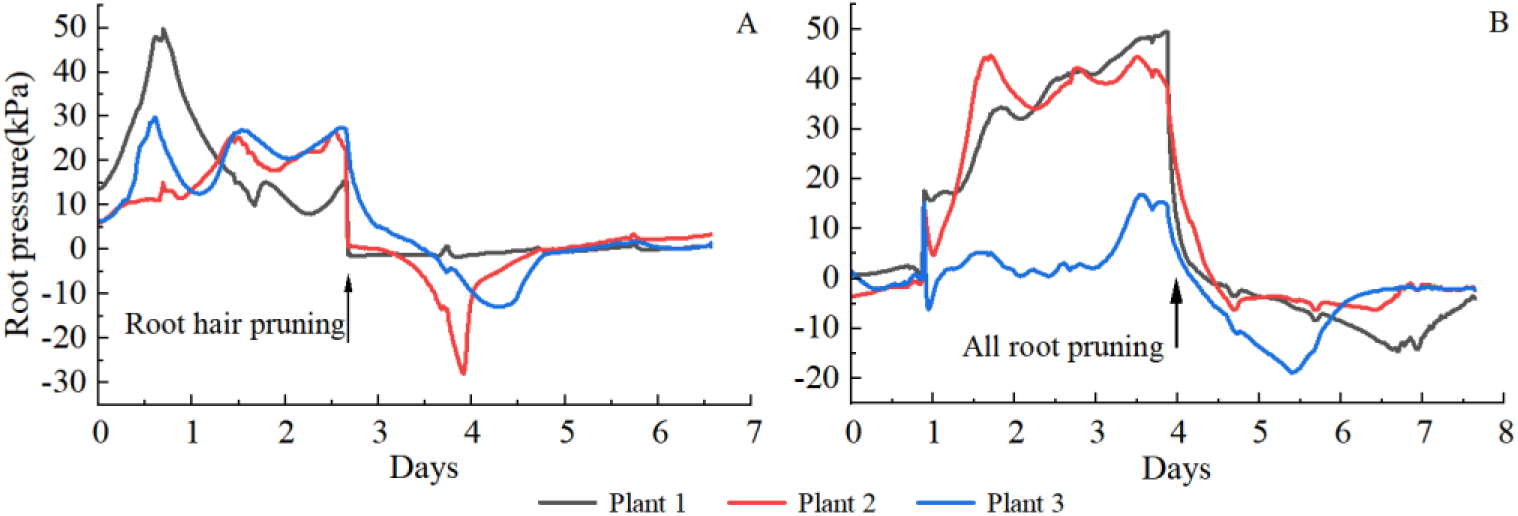
Root pressure of *D. sanderiana* after cutting off the fibrous roots.

## Discussions

*D. sanderiana* consistently exhibited significant positive root pressure throughout the day, reaching a peak value of 80 kPa, which is adequate to transport water up to 8 meters, despite the plants not exceeding 1 meter in height. In comparison, a measured root pressure of 64 kPa near the base of the stem in *Doliocarpus brevipedicellatus*, a species capable of reaching heights of 18 meters, can only push water up to approximately 6.4 meters (Ewers *et al*., 1997). Above this height, the pressure in the xylem approaches zero (Ewers *et al*., 1997). Furthermore, a root pressure of 64 kPa is insufficient to fully repair embolism at the tops of trees measuring 7.1 meters in height (Singh, 2016). The consistent positive root pressure exhibited by *D. sanderiana* renders it an exemplary candidate for root pressure studies. This can eliminate the influence of transpiration pull in the cohesion-tension theory (C-T theory). The lower or even negative root pressures observed in *D. sanderiana* may be associated with a decline in root vitality or closure of wounds due to prolonged measurement. The root pressure of *D. sanderiana* displayed a significant diurnal rhythm, with higher levels during the day and lower levels at night, in accordance with numerous previous studies (Grossenbacher, 1938; Parsons & Kramer, 1974; Guo & Wan, 2017). The variation in root pressure may not be solely attributed to the diurnal temperature fluctuations present in the laboratory, but could also be associated with the photoperiod established during the growth (Parsons & Kramer, 1974).

The generation and magnitude of root pressure are also influenced by soil moisture (Singh, 2016; Meunier *et al*., 2017). Gymnosperms and angiosperms grown on dry soil typically do not exhibit root pressure, whereas trees in moist areas demonstrate noticeable root pressure (White *et al*., 1958). Under drought stress conditions (soil moisture content below 45%), plants such as *Plectranthus scutellarioides*, sunflowers (*Helianthus annuus*), and tomatoes (*Solanum lycopersicum L*.) cultivated in sandy soil cease the production of root pressure, but resume production after rehydration (Jackson *et al*., 1996; Emerman & Dawson, 1997). Generally, plants exhibit higher root pressure when adequately hydrated, as evidenced in *Rhipidocladum racemiflorum*, where root pressure immediately increases following rainfall (Cochard, H. *et al*., 1994). The stronger the drought stress, the greater the rate of decline in root pressure in *D. sanderiana*. Consequently, there may be significant differences in root pressure between hydroponic and soil-cultured plants, with hydroponic plants potentially demonstrating greater root pressure due to availability to sufficient water. Measurements of root pressure in soil-cultured lucky bamboo at field capacity revealed similar magnitudes to those observed in hydroponic condition.

NaCl weakens the water-absorbing capability of roots, leading to a reduction in root pressure and eventual dehydration-induced plant death (Zhou, *et al*., 2023). Under low concentrations of NaCl (1 and 10 mmol·L^-1^), the root pressure of *D. sanderiana* initially increases, indicating that at first, NaCl acted as a nutrient promoting plant growth, given that the previous solution was plain water with low salt content. It was only after a period of accumulation that the toxic effects of NaCl became evident. In contrast, at high concentrations (100 and 200 mmol·L^-1^), salt immediately exerts toxic effects on plants due to the excessive absorption of NaCl by the plant. Research on *Suaeda salsa* suggests that, at appropriate NaCl concentrations, plants can alleviate osmotic stress by absorbing Na^+^ ions for osmoregulation, maintaining cell turgor, and mitigating the impact of osmotic stress on growth and photosynthesis (Bai *et al*., 2012). However, excessive NaCl exacerbates water stress and is detrimental to plants (Bai *et al*., 2012). Sodium chloride stress can affect plant growth through both direct and indirect mechanisms. Its direct effects primarily involve reducing soil water potential, impeding plant root water uptake, and inducing toxicity due to excessive accumulation of salt ions within the plant (Luo *et al*., 2003). Root pressure plays a pivotal role in evaluating the beneficial and detrimental effects of various mineral elements and their concentrations on plants. It can promptly and intuitively indicate these effects without necessitating visible damage to occur to the plants. In future studies, we may explore the impacts of various ions on plants using root pressure.

The magnitude and peak value of root pressure exhibit variation in response to temperature, increasing with rising temperatures and decreasing conversely (Zimmermann, 19 83; Ewers *et al*., 2001; Guo & Wan, 2017; Singh, 2016). This study also supports this conclusion. Root pressure is an energy-dependent process; enzymes involved in root activity are regulated by temperature, influencing root respiration metabolism and water transport (Ewers *et al*., 2001; Zhang *et al*., 2022). Within the physiological range of temperatures, for every 10℃ increase in temperature, plant respiration rate doubles to quadruples (Parsons & Kramer, 1974). However, our study found that at a temperature of 40℃, there was no observed increase in root pressure but instead a rapid decrease. This suggests that a water temperature of 40℃ caused damage to *D. sanderiana* plants leading to reduced vitality and subsequently lower root pressure. Additionally, it has been reported that low temperatures significantly alter the diurnal rhythm of root pressure although the specific mechanism remains unclear (Jelena *et al*., 2013). Concerning whether root pressure still exhibits diurnal rhythm under constant temperature conditions, Grossenbacher (1938) observed that the root pressure of hydroponically cultivated plants exhibits periodic variations under constant environmental conditions, peaking in the afternoon and reaching its lowest point in the morning. The diurnal cycle of root pressure may be determined by physiological rhythms established during growth, rather than being directly influenced by temperature and light cycles at that time (Grossenbacher, 1938; Parsons and Kramer, 1974) observed t hat midday stem bleeding in plants is 2–3 times higher than at midnight, and after eight days of continuous illumination, the diurnal rhythm of root pressure disappears (Parsons & Kramer, 1974), indicating the transmission of signaling substances (such as sugars or growth regulators) from leaves to roots. The substances secreted by leaves regulate the synthesis and distribution of root AQP to modulate active water absorption by roots (Javot & Maurel, 2002).

Mineral elements play a crucial role in influencing root pressure. For instance, in walnuts, when the soil nitrate concentration decreased, the xylem pressure ranged between 6 0–100 kPa; conversely, an increase in nitrate concentration led to a rise in xylem pressure to 130 kPa within 4 hours (Ewers *et al*., 2001). This phenomenon may be attributed to th e active absorption mechanism of mineral elements by plant root systems. The uptake of mineral elements by plant roots generates an osmotic potential, inducing directional water flow and resulting in the development of root pressure (Ewers *et al*., 2001). However, this study observed a decrease in root pressure when the nitrate concentration reached 4 g·L^-1^. Additionally, Zhao *et al*. 2021 found that exceeding potassium nitrate concentrations o f 30 mmol·L^-1^ (3 g·L^-1^) resulted in reduced radicle length and fresh weight of *Brassica rapa var. glabra* Regel seedlings with increasing KNO3 concentrations. Furthermore, Gu *et al*. 2023 observed that when nitrate concentrations exceeded 50 mmol·L^-1^ (5 g·L^-1^), the enzymatic activity of POD and CAT in the leaves of Calathea decreased, impeding the elimination of reactive oxygen species, resulting in cell membrane damage and leading to slowed or halted plant growth (Gu *et al*., 2023). Therefore, it can be inferred that excessive nutrient stress causing cell membrane damage leads to diminished water transport capability and subsequent decline in root pressure when mineral element concentrations exceed 40 mmol·L^-1^(4 g·L^-1^), highlighting the need for further comprehensive investigation into the type and concentration of nutritional elements such as nitrates and their impact on root pressure. This will contribute to enhancing our understanding of the role of mineral elements in plant physiological processes.

The leaves located below the stem have a significant influence on root pressure. The presence of leaves beneath the stem leads to a near-zero level of root pressure. This occurrence is due to the direct interference of these lower stem leaves with water transport in the upper stem, causing root pressure to approach zero. Furthermore, the removal of leaves has an impact on root pressure. Root pressure slightly increases after all leaves are removed, but this does not alter the rhythmic changes in root pressure. Zhang *et al*., 2023 has demonstrated that tillering results in a reduction of root pressure (Zhang *et al*., 2023). This decrease may be attributed to the leaves at the tillering sites consuming water through transpiration, thereby diminishing the pressure experienced by the stem above. We deduce that the impact of tillering is akin to that of leaves. Even though root pressure values are minimal when leaves are present, the root pressure at the leaf location is actually above zero, indicating that root pressure functions fully if the pressure sensor is positioned above the leaves. Shi (2024) has indicated that stem bleeding phenomena in *Acer buergerianum* and *Acer×freemanii‘Sienna Glen’* occur from mid-November to mid-March of the following year, with no stem sap flow observed after full leaf expansion in April (Shi, 2024).

The removal of all fine roots or root systems from lucky bamboo has a significant impact on root pressure, causing it to become negative. A comparison between the effects of temperature (Fig. 4) and the removal of fine roots (Fig. 7) reveals that the decrease in water temperature has a more rapid impact on root pressure. After removing the root hairs, it takes several hours for the root pressure to decrease to 0, whereas low temperature results in an almost instantaneous and substantial decrease in root pressure. This indicates that metabolic physiological activities in the roots result in a rapid decrease in root pressure, which cannot be promptly and effectively regulated after the removal of root hairs. The removal of root hairs causes the plant to lose a significant area of water-absorbing tissue and disrupts the regulation of AQP. The role of AQPs and carrier proteins in root pressure generation is gaining increasing attention (Wegner, 2014; Fricke, 2015). AQPs are closely associated with root water absorption, as a large number of AQPs on the plasma membrane actively participate in water transport (Zhai *et al*., 2017). When AQP activity is inhibited with HgCl_2_, water conductivity decreases by 20%–90%. Additionally, AQPs function as pressure-driven gated channels (Wan *et al*., 2004; Chaumont & Tyerman, 2014). Johansson (1998) discovered that the gating mechanism of plant AQPs is regulated by turgor pressure, suggesting that turgor pressure may serve as a source of root pressure (Enns *et al*., 2000).

## Conclusion

The pressure sensor method is suitable for quantifying root pressure in herbaceous plants with relatively rigid stems. *D. sanderiana*, when grown in soil or hydroponic culture, consistently exhibits positive root pressure throughout the day, significantly surpassing the pressure required to support its water height. It exhibits a diurnal pattern with higher values during the day and lower values at night. This sustained positive root pressure enables us to eliminate the influence of transpirational pull and focus exclusively on studying root pressure. Root pressure decreases with increasing concentrations of PEG 6000, maintaining diurnal rhythm changes at low concentrations; however, at higher concentrations, it declines rapidly and without any diurnal rhythm. Low concentrations of NaCl temporarily lead to a significant increase in root pressure of *D. sanderiana*, which subsequently gradually decreases under the influence of NaCl. Conversely, high concentrations of NaCl swiftly diminish the plant’s vigor, resulting in a rapid decline in root pressure. Root pressure fluctuates with changes in temperature; it rises with increasing temperatures and declines with decreasing temperatures at room temperature, leading to alterations in the diurnal rhythm of root pressure. At 40℃, the root pressure of the plant experiences a rapid decline. Elevated nitrate concentrations lead to an increase in root pressure, but once a specific concentration is reached, root pressure diminishes while retaining diurnal rhythm variations, suggesting that the root pressure is influenced by the mineral element content. *D. sanderiana* with leaves positioned below the pressure measurement point demonstrates lower root pressure, akin to the majority of other plants during root pressure measurements. The removal of fibrous roots and the entire root system also exerts an influence on root pressure, leading to a rapid decrease. Water, temperature, nitrates, leaves, as well as the removal of fibrous roots and the entire root system have definitive impacts on root pressure, underscoring the essential role of environmental factors in measuring root pressure. The use of hydroponic *D. sanderiana* provides the advantages of reusability and enables continuous, precise control over environmental factors such as temperature, drought, nutrients, and salinity gradients. This approach offers a more robust scientific method for investigating root pressure and its underlying mechanisms.

